# Network-guided identification of candidate biomarkers for COPD-associated comorbidities using deep learning and real-world transcriptomic data

**DOI:** 10.64898/2026.10.26.761000

**Authors:** Jose Manuel Mas, Esther Ramírez, Xavier Daura

## Abstract

Chronic obstructive pulmonary disease (COPD) frequently coexists with infections, metabolic disease, cardiovascular disease, inflammatory lung injury and lung cancer. Molecular markers derived by comparing a comorbidity with healthy controls need not identify the transition from an established COPD state to that comorbidity. We therefore developed a two-stage computational strategy that asks a narrower question: which proteins can distinguish COPD samples from samples annotated with a specific COPD-associated comorbidity?

First, a Human Protein Network (HPN; 17,302 proteins and 492,195 relationships) was encoded by 20 graph-derived features. A convolutional-neural-network ensemble, termed the functional-distance model (FDM), was trained on known drug-target–indication relationships. Stored training-set predictions yielded an apparent balanced accuracy of 0.811 and Matthews correlation coefficient (MCC) of 0.627; these values describe resubstitution performance and are not an independent estimate of generalization. The COPD-centred network search evaluated 2,645 nearby proteins and retained 224 candidates using a COPD FDM score above 0.50. Candidate expression was then tested in independently assembled public patient-derived transcriptomic data using simple one-feature classifiers and cross-validation.

The resulting exploratory panel contained GTF2I for bacterial infection, UBXN7 for type 2 diabetes, LIFR for hypertension, SRGN for lung neoplasm and FGF4 for pneumonitis. Across the saved cross-validation predictions, balanced accuracy ranged from 0.838 to 0.970 and MCC from 0.726 to 0.941. SRGN has direct experimental support in non-small-cell lung cancer, whereas the remaining candidates have varying indirect mechanistic support and should be regarded as hypotheses rather than established disease biomarkers.

This work demonstrates an orthogonal discovery framework in which network-based functional prioritization and patient-derived molecular discrimination constrain one another. Its main contribution is methodological: it provides a reusable strategy for generating comorbidity-specific biomarker hypotheses from an index-disease state. Prospective longitudinal cohorts are required before claims of progression prediction or clinical utility can be made.

## Introduction

COPD is a heterogeneous chronic respiratory disease and a major global cause of morbidity and mortality [1–3]. Its clinical course is shaped not only by airflow limitation but also by acute infections and by metabolic, cardiovascular, inflammatory and malignant comorbidities [4,5]. These conditions influence symptoms, treatment decisions, hospitalization and survival, yet they arise against a molecular background already altered by COPD, smoking exposure, ageing and systemic inflammation.

Early recognition of a comorbidity can improve clinical surveillance and patient stratification, but molecular markers are commonly discovered by comparing a single disease with healthy controls. That design may identify markers of the comorbidity itself while overlooking signals that remain informative when the reference state is already COPD. A biomarker intended to support surveillance of COPD-associated comorbidity should therefore answer a different question: does the molecular signal distinguish an established COPD state from the state represented by a specific associated condition?

Computational and systems-biology approaches can generate such hypotheses at a scale that would be impractical experimentally. However, their false-positive rate and the cost of downstream clinical validation make an independent evidence stream essential. Patient repositories containing both molecular measurements and interpretable clinical metadata are particularly valuable for this purpose, but open repositories assembled explicitly for biomarker validation remain uncommon.

Gene Expression Omnibus (GEO) was not created as a clinical patient repository; it is a public archive of functional-genomics studies [6]. Nevertheless, a substantial fraction of its records correspond to human patients and include disease, tissue and study metadata. The project therefore treated GEO as a source from which a patient-derived real-world molecular repository could be constructed after systematic metadata curation, pathology harmonization and within-study transformation. In the project-wide extraction, millions of human-labelled records were screened and nearly two million samples were considered sufficiently patient-oriented for downstream curation across 314 pathology labels. These figures describe the repository-building programme, not the effective sample size of any individual classifier.

This reuse of public transcriptomics extends the idea of real-world data (RWD) beyond routinely collected electronic health records. The measurements were generated for heterogeneous research purposes rather than under one prospective protocol, so they cannot be treated as a conventional clinical cohort. Their value lies instead in providing patient-derived molecular observations, preserved with study provenance, that can test predictions generated independently by network biology [7]. The same repository concept can support future studies of other diseases, drug effects and comorbidity systems.

Systems biology supplies the complementary evidence stream. A Human Protein Network (HPN) represents proteins as nodes and curated physical or functional relationships as edges. Graph topology summarizes proximity, overlap and connectivity between candidate proteins and disease effectors, but topology alone can generate nonspecific associations. Combining network-based prioritization with patient-derived expression therefore creates an orthogonal filter: the HPN proposes functionally plausible candidates, and RWD tests whether those candidates discriminate the intended clinical contrast.

Here we apply this framework to bacterial infection, type 2 diabetes, hypertension, lung neoplasm and pneumonitis in the context of COPD. Our objective is not to identify biomarkers of COPD itself. It is to generate candidate proteins that distinguish COPD samples from samples representing selected COPD-associated comorbidities, while documenting the reusable methodology for constructing the HPN, training the functional-distance model and converting GEO into a patient-derived molecular resource. Cross-sectional discrimination is reported without claiming temporal prediction, causal involvement or immediate clinical utility. Importantly, the study does not rely on samples from patients simultaneously diagnosed with COPD and the corresponding comorbidity. Instead, it compares independent patient cohorts representing COPD and each selected comorbidity. This approach aims to identify molecular features that distinguish comorbidity-associated disease states from the molecular background of COPD, generating candidate biomarkers for subsequent evaluation in patients with both conditions.

## Methodology and Data Sources

### Overall strategy

The workflow combines two independent computational stages. The first stage uses an HPN and a deep-learning functional-distance model (FDM) to identify proteins with topological patterns compatible with a functional relationship to COPD. The second stage evaluates those candidates in patient-derived expression data. A protein reaches the final panel only when it is supported by the network-based screen and can separate COPD from one selected comorbidity using a transparent one-feature model. This separation between hypothesis generation and molecular testing is central to the design (Figure 1).

**Figure 1.**
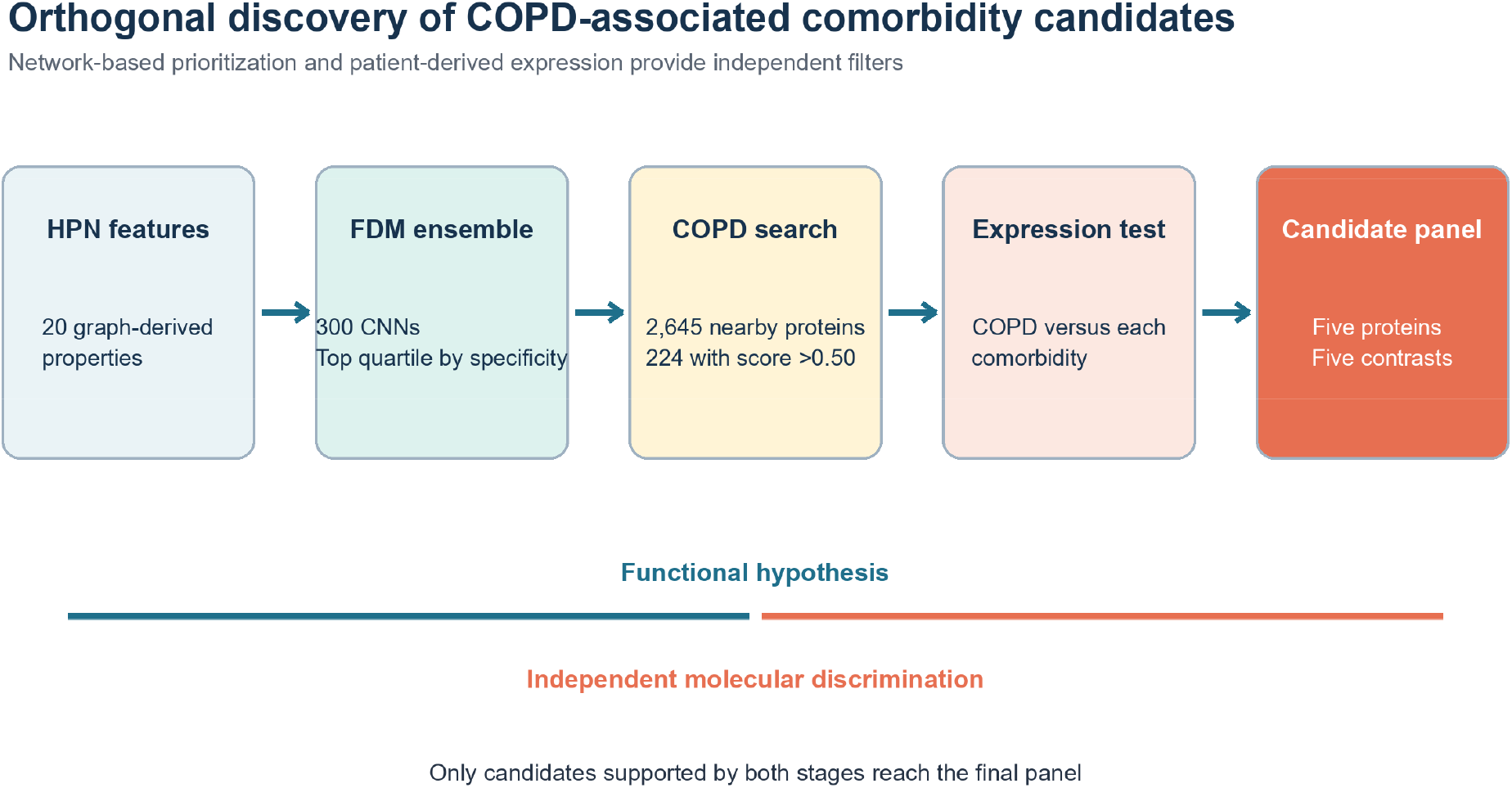
Orthogonal strategy for discovering COPD-associated comorbidity candidates. Network topology generates functional hypotheses; an independently assembled patient-derived expression repository tests molecular discrimination. The final panel contains one candidate for each analysed COPD-comorbidity contrast.

### Preparing GEO as a patient-derived RWD repository

GEO series and samples were screened for human-derived expression data and informative metadata. Cell lines, experimental systems without a patient interpretation, poorly documented samples and records without a usable molecular matrix were excluded from the patient-oriented layer. Free-text disease descriptions, study titles, sample characteristics and tissue annotations were combined to map samples to a controlled pathology vocabulary. The resulting repository retained the original study and sample identifiers so that every analytical record remained traceable to its source (Figure 2).

**Figure 2.**
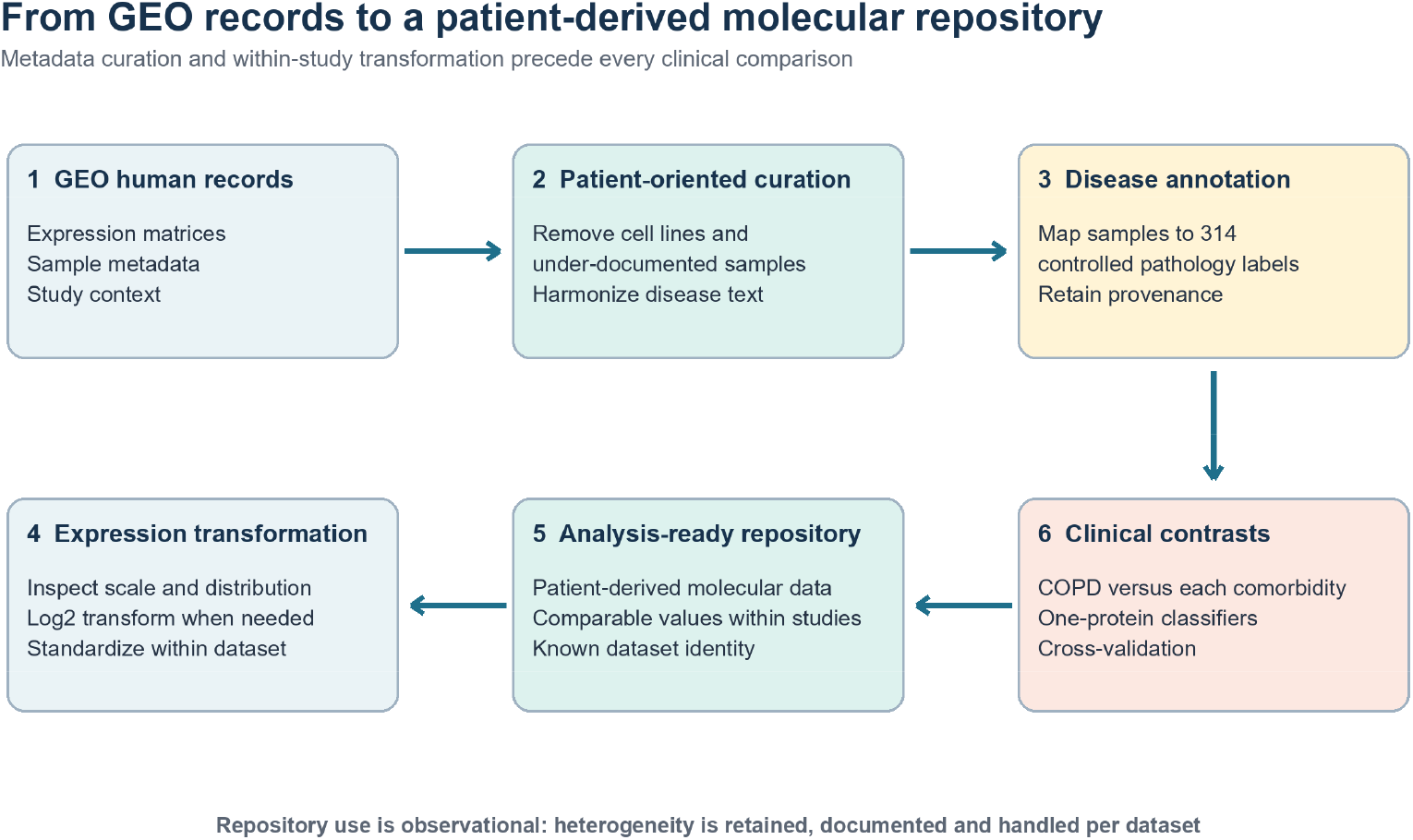
Construction of the patient-derived molecular repository from GEO. Metadata curation, pathology harmonization and within-study transformation convert public transcriptomic records into traceable analysis-ready observations. Repository-scale counts and classifier-specific effective sample sizes are deliberately kept separate.

The repository does not imply that all GEO samples are mutually exchangeable. Measurements differ by platform, tissue, preprocessing and study design. Transformations were therefore performed within the relevant dataset rather than by applying a single global normalization across the entire archive. If a measurement vector x was already approximately log2-scaled or normally distributed, it was standardized as

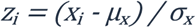

If the distribution indicated an untransformed positive scale, a small constant ε was added before log2 transformation and the transformed values y were then standardized:

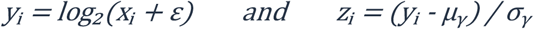

This protocol places values on a comparable standardized scale within each source dataset while preserving the distinction between studies. It reduces scale-driven artefacts but does not remove biological or technical batch effects. Consequently, the repository is best understood as an observational, patient-derived molecular resource for computational validation, not as a prospectively enrolled or uniformly processed clinical cohort (Figure 3).

**Figure 3.**
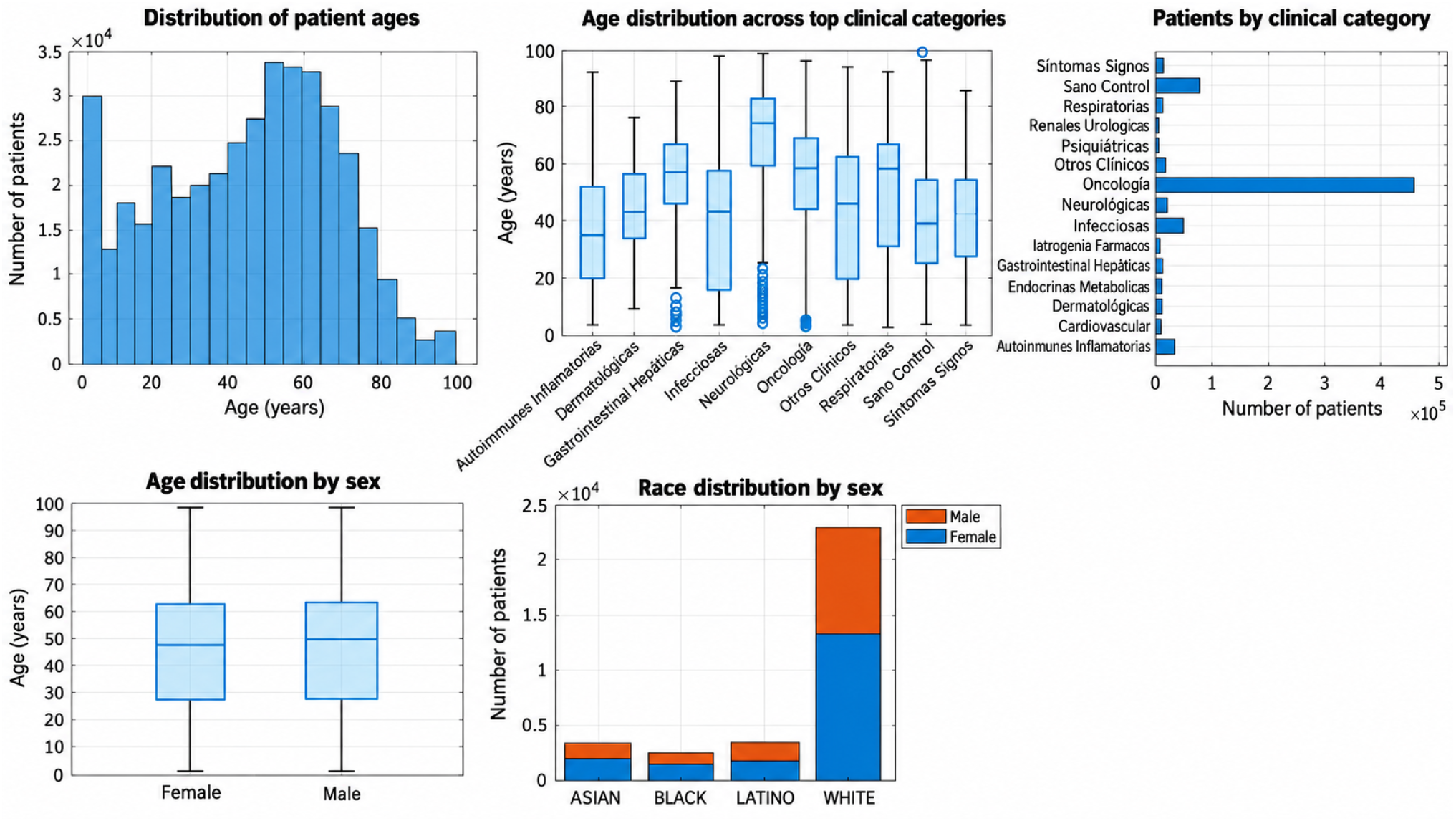
Several distributions with information about the number of patients, age, sex, race and types of pathologies describes the content of our patient repository of RWD ready to be exploited.

### Repository coverage for the analysed disease categories

After curation, the archived repository inventory contained the sample counts shown below. These values describe disease-labelled records potentially available before protein-specific missingness, study eligibility and classifier-specific filtering. They are therefore repository-coverage figures rather than the effective sample sizes of the final models, which are reported separately in Table 3. Pulmonary arterial hypertension and systemic hypertension were treated as distinct clinical entities. Pulmonary arterial hypertension was represented in the archived repository inventory and HPN disease-effector annotations (Tables 1 and 2), whereas systemic hypertension was used for the transcriptomic biomarker discrimination analysis. No patients annotated with pulmonary arterial hypertension were available for that analysis. Accordingly, the LIFR findings refer specifically to systemic hypertension and should not be extrapolated to pulmonary arterial hypertension.

**Table 1.** Archived RWD repository coverage for the disease categories considered in the study. Counts characterize the available curated records and must not be interpreted as mutually independent observations or as classifier-specific sample sizes.

| Repository category | Available samples |
| --- | --- |
| Healthy reference records | 3,799 |
| Bacterial infection | 754 |
| Type 2 diabetes | 184 |
| Lung neoplasm | 312 |
| Chronic obstructive pulmonary disease | 3,275 |
| Pneumonitis | 87 |
| Hypertension | 199 |

**Table 2.** Numbers of curated mediator or effector proteins available in the HPN for COPD and the archived comorbidity categories.

| HPN pathology category | Mediator or effector proteins |
| --- | --- |
| Bacterial infection | 72 |
| Type 2 diabetes | 175 |
| Lung neoplasm | 104 |
| Chronic obstructive pulmonary disease | 88 |
| Pneumonitis | 170 |
| Pulmonary arterial hypertension | 50 |

**Table 3.** Final five-protein exploratory panel. Effective n and metrics were reconstructed from the saved cross-validation confusion matrices. *F1 is positive-class dependent and is retained for traceability to the original selection procedure.

| COPD contrast | Candidate | One-feature model | Effective n | Balanced accuracy | MCC | F1* |
| --- | --- | --- | --- | --- | --- | --- |
| Bacterial infection | GTF2I<br>P78347 | Linear regression + threshold | 400 | 0.970 | 0.941 | 0.969 |
| Type 2 diabetes | UBXN7<br>O94888 | Optimized quadratic threshold | 311 | 0.883 | 0.788 | 0.927 |
| Hypertension | LIFR<br>P42702 | Optimized quadratic threshold | 400 | 0.897 | 0.806 | 0.888 |
| Lung neoplasm | SRGN<br>P10124 | Optimized threshold | 359 | 0.838 | 0.726 | 0.884 |
| Pneumonitis | FGF4<br>P08620 | Linear regression + threshold | 273 | 0.963 | 0.925 | 0.980 |

### Construction of the Human Protein Network

The HPN represents human proteins as nodes and experimentally or knowledge-supported relationships as edges. Its purpose is not to reproduce a single protein-protein interaction database, but to combine complementary biological layers into a common UniProt-centred graph. KEGG and Reactome contributed pathway, reaction and metabolic relationships; IntAct, BioGRID and HPRD contributed curated physical and genetic interactions; and TRRUST contributed transcription-factor and target-gene regulation [8–13]. Relations were typed according to their source evidence, identifiers were harmonized, duplicate links and self-relations were controlled, and provenance was retained during integration.

The graph was then enriched with disease effector proteins, drug targets and their approved or documented indications, tissue-expression information and known biomarker annotations. Disease effectors were assigned through the project knowledge base and literature curation, whereas drug-target and indication information drew on DrugBank and PubChem [14,15]. These annotations provide the functional labels used for FDM training and later permit the same HPN to support drug-repurposing, adverse-event and disease-mechanism analyses.

Network hubs can create artificially short paths between otherwise unrelated protein sets. Proteins connected to more than 300 neighbours were therefore treated as sticky proteins and excluded from the distance calculations used here. This filtering does not assert that the proteins are biologically unimportant; it prevents highly promiscuous connectors from dominating topological similarity. The resulting HPN version contains 17,302 proteins and 492,195 relationships, together with effector annotations for 314 pathologies and 3,924 drug targets (Figure 4).

**Figure 4.**
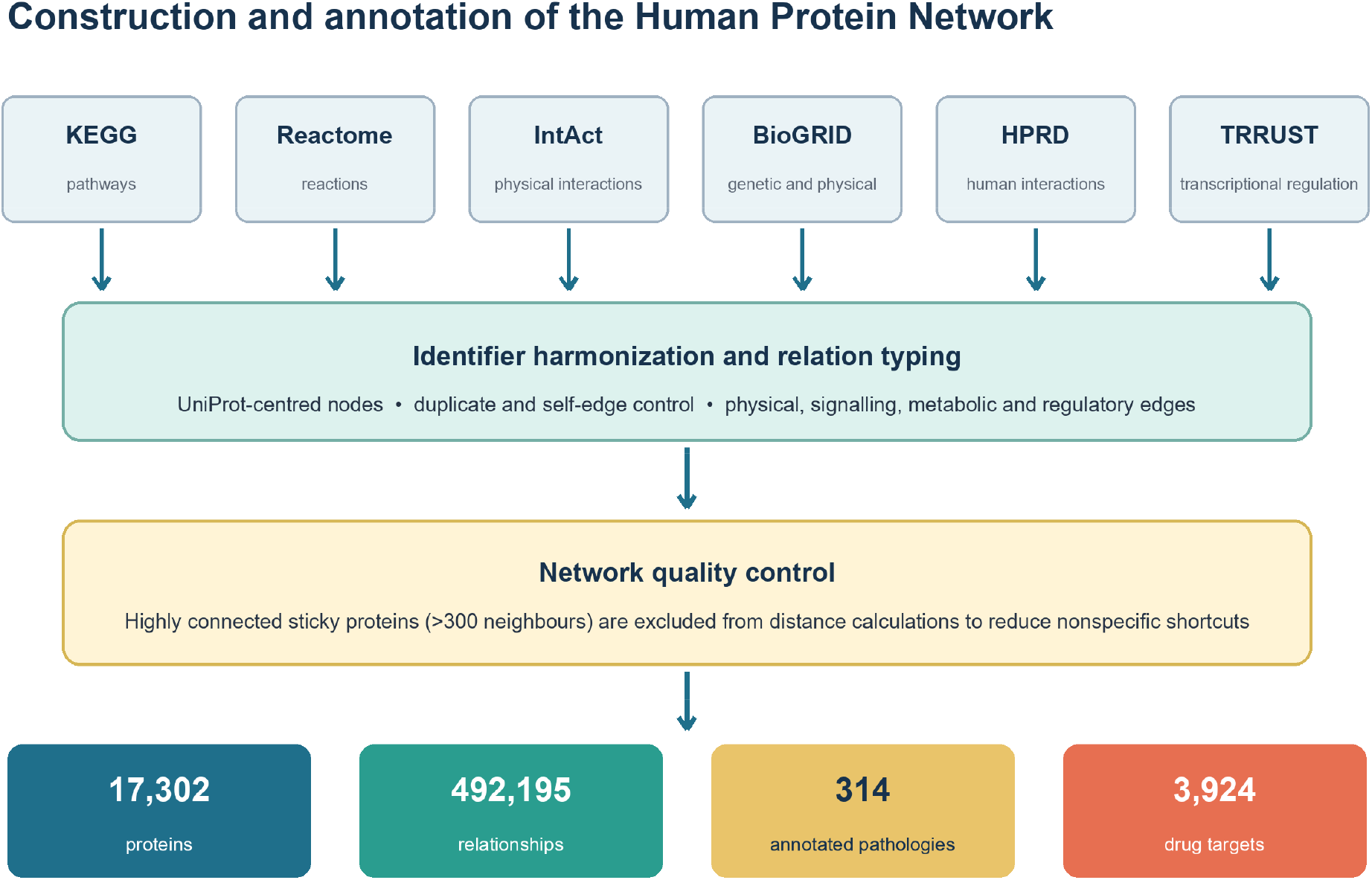
Construction and annotation of the HPN. Complementary pathway, interaction and regulatory resources are harmonized in a common protein graph. Sticky proteins with more than 300 neighbours are excluded from distance calculations to reduce nonspecific network shortcuts.

### Disease-effector coverage in the HPN

The pathology annotations used to define disease-centred protein sets were not uniform in size. The numbers of curated mediator or effector proteins available for the archived COPD and comorbidity categories are reported below. These counts document the network evidence available to the distance calculations; they do not measure disease prevalence or biomarker performance.

### Functional-distance representation and model training

The FDM was trained to recognize topological configurations associated with known functional relationships. For each drug target and pathology-effector set, 20 graph-derived properties were calculated. They included shortest-path summaries, average distances, Hausdorff- and Frechet-type set distances, Jaccard similarity and distance, Dijkstra-derived paths and related neighbourhood descriptors. Known target-indication pairs were assigned to one class and target-non-indication pairs to the other. This labelling turns curated pharmacological knowledge into a general training signal for functional proximity in the HPN.

The source matrix contained 554,817 records with 20 features: 227,240 positive and 327,577 negative labels. Balancing produced 655,154 training records. Because neighbouring graph measurements are not fully independent, an ensemble of 100 random forests was used to order the features before CNN training. The implemented CNN receives a 20 × 1 × 1 tensor and applies two convolutional/ReLU/pooling blocks along the ordered feature axis, followed by a fully connected classification head. The saved architecture contains no dropout layer.

Three hundred CNNs were trained using stochastic gradient descent with momentum, an initial learning rate of 0.01 and up to 20 epochs. The top quartile by specificity formed the FDM metamodel. Its aggregate values in this paper were recalculated from stored training-set predictions and are therefore apparent or resubstitution estimates. They justify use as a prioritization score within the workflow but do not constitute an independent estimate of generalization.

### Candidate generation and RWD validation

The COPD-centred search considered proteins in the predefined local HPN neighbourhood and its one-step expansion. This produced 2,645 proteins for scoring. Proteins with an aggregate COPD FDM score above 0.50 were retained, yielding 224 candidates. This definition deliberately uses only the reproducible COPD score; legacy comorbidity-specific CNN columns were excluded because their historical label handling cannot be reconstructed with sufficient confidence. For each clinical contrast, transcriptomic samples annotated with the selected comorbidity were compared against samples annotated with COPD. These were independent disease cohorts rather than patients with confirmed concurrent diagnoses. Consequently, the resulting classifiers identify molecular differences between disease states and should be interpreted as exploratory candidates for COPD-associated comorbidity discrimination, not as validated markers of comorbidity onset in COPD patients. Comparisons against healthy controls were not used to establish that the identified signals were exclusive to the COPD–comorbidity contrast.

For each candidate and COPD-comorbidity contrast, simple one-feature classifiers were evaluated: a single optimized threshold, a two-threshold or quadratic rule for non-monotonic separation, and a linear-regression-based separator. The historical final panel selected the highest saved cross-validated F1 candidate for each contrast. In the revised report, balanced accuracy and Matthews correlation coefficient (MCC) are primary because they remain unchanged when the binary labels are exchanged:

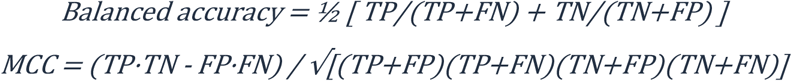

F1 is retained as a secondary traceability measure because it determined the historical ranking, but it depends on which class was encoded as positive. Effective sample sizes were reconstructed directly from the stored predictions as TP + TN + FP + FN. No external holdout, multiplicity-adjusted inference, calibration analysis or prospective risk estimate was available in the archived outputs.

### Literature contextualization

Each selected protein was searched by approved symbol and UniProt identifier together with COPD, the corresponding comorbidity and mechanistically adjacent terms. Primary experimental or patient-based studies were prioritized. Evidence was classified as direct only when the same protein was investigated in the relevant disease family. Pathway, ligand-receptor, animal-model, stress-response and gene-locus associations were classified as indirect. Evidence involving circUBXN7 was explicitly kept separate from evidence for the UBXN7 protein.

## Results

### The repository and HPN provide complementary coverage of the selected conditions

The curated RWD inventory contained between 87 pneumonitis records and 3,799 healthy reference records across the archived categories, together with 3,275 COPD records (Table 1). The HPN annotation layer contained between 50 and 175 mediator or effector proteins per selected condition (Table 2). These two coverage measures describe different resources: repository counts indicate potentially available patient-derived observations, whereas effector counts define the network sets used for functional-distance calculations. Neither substitutes for the effective sample size of a protein-specific classifier.

### The FDM provides a reproducible prioritization score, with performance caveats

The training matrix contained 554,817 original protein-pair records with 20 features (227,240 positive and 327,577 negative labels); class balancing produced 655,154 training records. The implemented network is formally a two-dimensional CNN receiving a 20 × 1 × 1 tensor. Because one spatial dimension has width one, the convolutions operate along the ordered feature sequence. Inspection of the stored model confirmed that it did not contain the 50% dropout layer described in the earlier manuscript. The ensemble comprised 300 CNN models; the top quartile was selected by specificity to form the FDM metamodel.

Aggregate metrics recalculated from stored predictions were balanced accuracy 0.8108, specificity 0.8732, precision 0.8551, recall/sensitivity 0.7484, positive-class F1 0.7982 and MCC 0.6266 (Figure 5). Balanced accuracy and MCC are emphasized because they are less sensitive to the arbitrary naming of the positive class than precision, recall or F1. Importantly, these values were obtained from predictions on the training records and therefore represent apparent or resubstitution performance. They support use of the FDM as a ranking instrument within this workflow, but not a claim of independently validated predictive accuracy.

**Figure 5.**
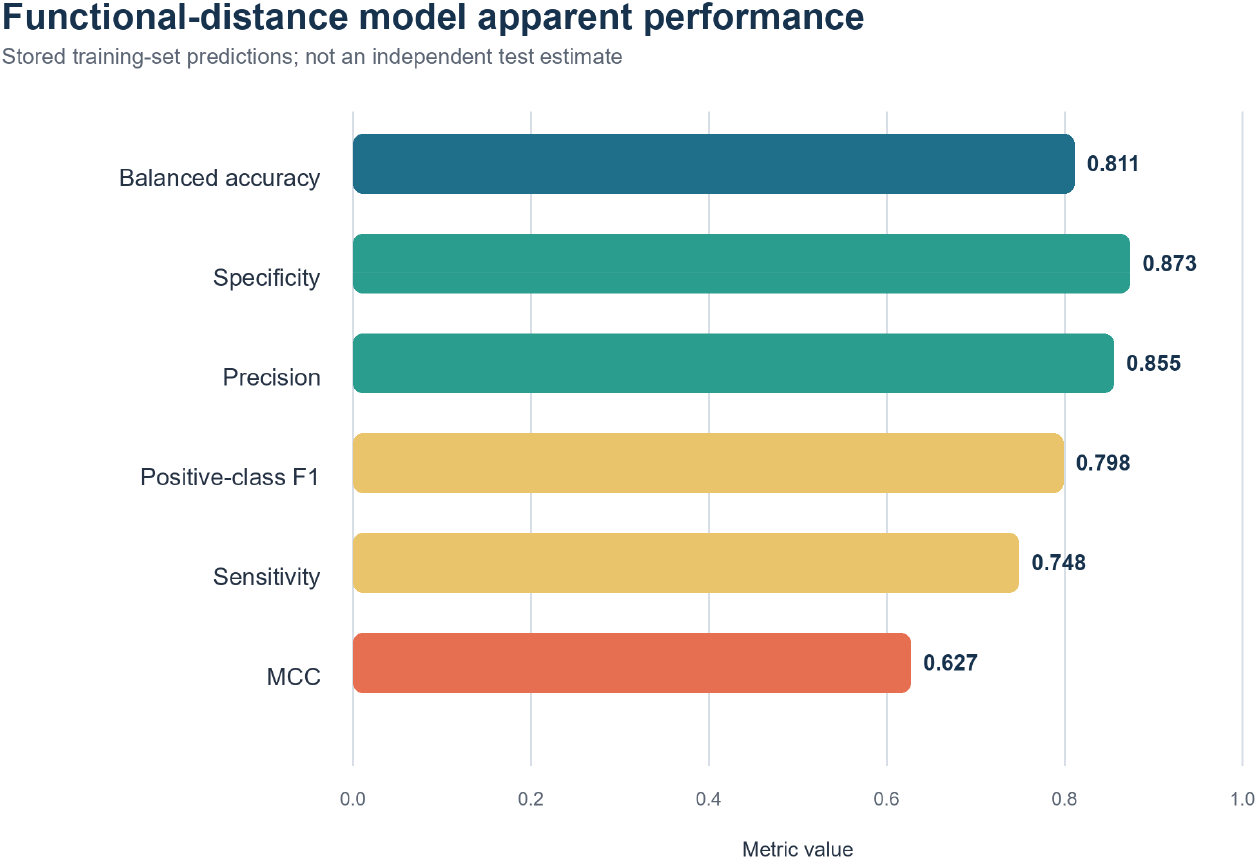
Aggregate FDM performance reconstructed from stored training-set predictions. Balanced accuracy and MCC are the primary summary measures. Precision, sensitivity and F1 depend on the selected positive class. All measures are apparent/resubstitution estimates and should not be interpreted as independent test performance.

### A COPD-centred network search yields 224 candidates

The FDM was applied to 2,645 proteins located in the COPD-centred network neighbourhood. Proteins with a COPD FDM score above 0.50 were retained, resulting in 224 candidates. This historical set remains reproducible because its definition uses only the COPD score. We deliberately did not retain earlier interpretations based on comorbidity-specific CNN output columns: a defect in the legacy prediction routine could assign labels to the wrong comparison, so those columns cannot support claims about direction or comorbidity-specific functional probability. The corrected routine is available for future runs, but the present manuscript does not retroactively reinterpret the affected values.

The archived expression screen also contained broader condition-specific lists of proteins meeting the historical rounded cross-validated F1 threshold of at least 0.90. Those lists demonstrate that the five reported proteins arose from a wider empirical screen rather than from direct nomination. Because multiplicity-adjusted inference and fully consistent provenance for every historical comparison label were not available, the wider lists are retained as exploratory project outputs and are not described as validated biomarkers in the main panel.

### Five one-protein models discriminate specific COPD–comorbidity contrasts

Each of the 224 proteins was evaluated using the stored patient-derived expression comparisons. The final panel selected the protein with the highest saved cross-validated F1 for each contrast; we retain that historical selection rule for traceability, while reporting balanced accuracy and MCC as the primary performance measures. The five selected models had balanced accuracy values of 0.838–0.970 and MCC values of 0.726– 0.941 (Figure 6; Table 3). These results concern discrimination between the two analysed cohorts; they do not establish whether expression is higher or lower in a clinically invariant direction, nor do they show that a marker predicts future comorbidity onset.

**Figure 6.**
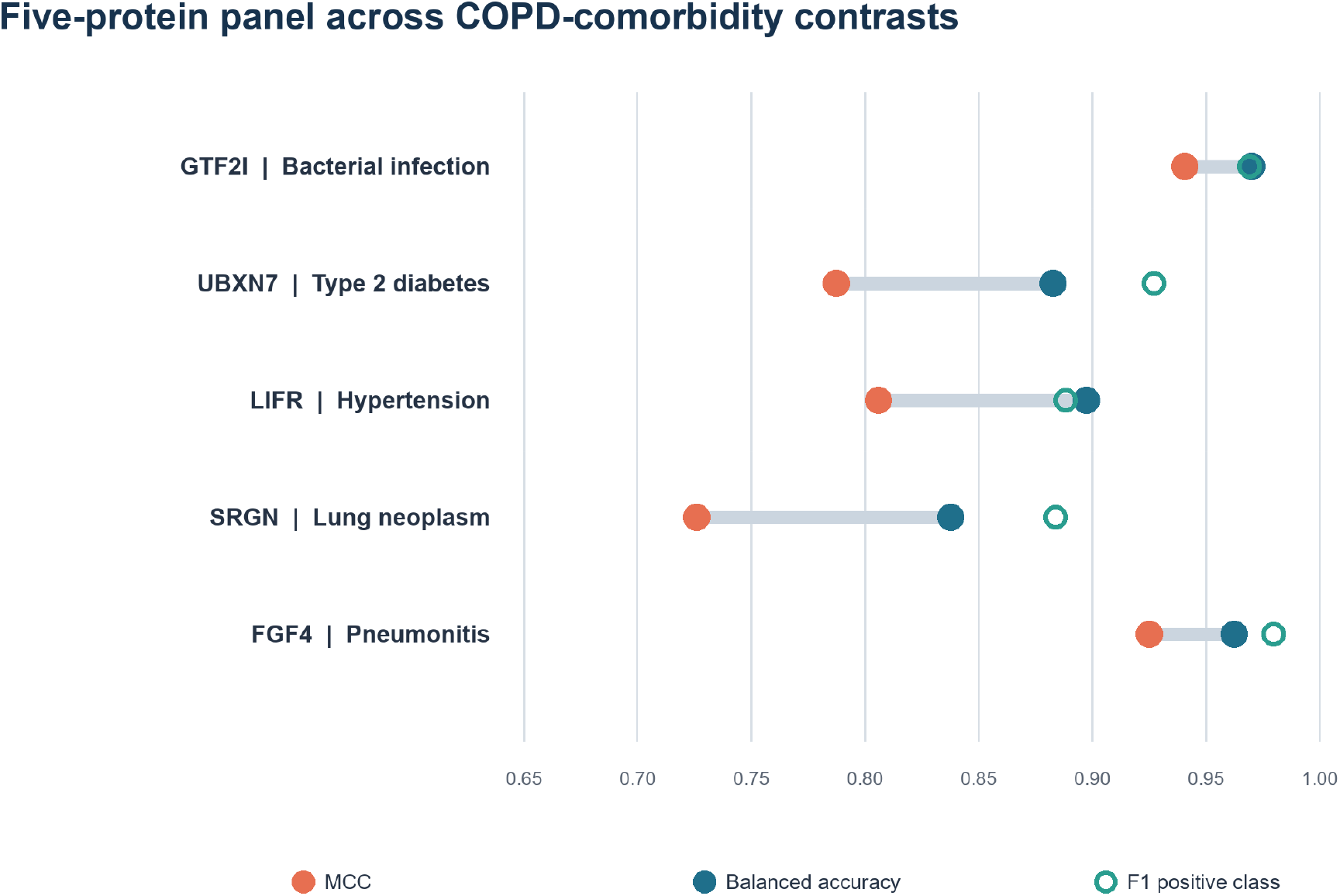
Cross-validated performance of the selected one-protein models. Balanced accuracy and MCC are label-symmetric primary metrics. F1 is included only to document the historical selection criterion and refers to the positive class stored in the analysis output.

The full saved confusion matrices are shown in Figure 7. Bacterial infection/GTF2I and pneumonitis/FGF4 displayed the most balanced class-wise results. The lung-neoplasm/SRGN model showed high sensitivity for the stored positive class but more false positives in the opposite class, which is reflected by its lower balanced accuracy and MCC. Reporting the matrices prevents a single summary score from obscuring class-specific error structure.

**Figure 7.**
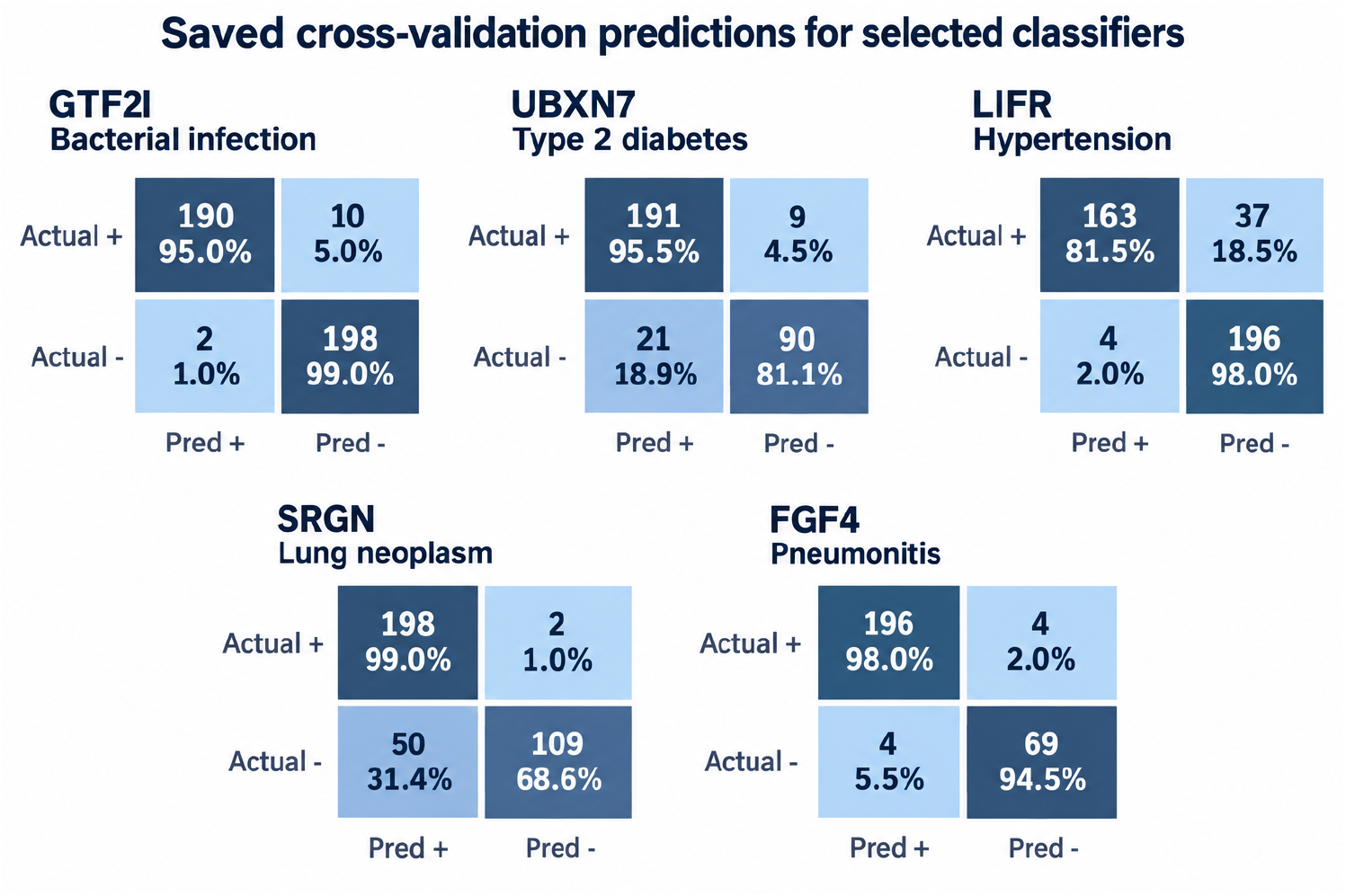
Confusion matrices from the saved cross-validation predictions. Counts and row percentages are shown. The signs indicate the positive and negative classes as stored in the historical analysis; the biological conclusions do not rely on interpreting one direction as increased or decreased expression.

### Literature context supports one candidate directly and four indirectly

The literature search was interpreted conservatively (Table 4). SRGN has the clearest external support: experimental studies report SRGN overexpression in primary non-small-cell lung cancers and demonstrate CD44-dependent migration, invasion, stemness and metastatic phenotypes [21,22]. This is direct disease-relevant evidence, although it does not independently validate the precise COPD-versus-lung-neoplasm classifier used here.

**Table 4.** Conservative literature interpretation. ‘Direct’ indicates experimental evidence involving the same protein and disease family; it does not denote clinical validation of the present model. ‘Indirect’ denotes pathway, model-system or locus-level support.

| Protein | Contrast | Evidence level | External biological context | Refs |
| --- | --- | --- | --- | --- |
| GTF2I | Bacterial infection | Indirect | Innate B-cell homeostasis and LPS-responsive transcription | [28] |
| UBXN7 | Type 2 diabetes | Indirect | NRF2/HIF-1α stress and metabolic-state regulation; circUBXN7 locus evidence is non-equivalent | [29,30] |
| LIFR | Hypertension | Indirect | COPD/smoke-related LIF signalling; cardiotrophin-1–LIFR axis in hypertensive remodelling | [23–26] |
| SRGN | Lung neoplasm | Direct | Overexpression and functional tumour-promoting activity in NSCLC | [21,22] |
| FGF4 | Pneumonitis | Indirect | Protection in LPS-induced acute lung injury and inflammatory signalling | [27] |

For LIFR, COPD-relevant data show reduced pulmonary LIF and LIFR signalling after cigarette-smoke exposure and in COPD-associated experimental contexts [23]. Hypertension support is mechanistic rather than marker-specific: cardiotrophin-1, which signals through the gp130/LIFR complex, is elevated in human hypertension and participates in cardiac, vascular and renal remodelling [24–26]. FGF4 attenuates lipopolysaccharide-induced acute lung injury through TLR4/NF-κB inhibition in animal and cellular models, providing lung-inflammation plausibility but not direct evidence in pneumonitis patients [27].

GTF2I regulates innate features of B lymphocytes and modulates LPS-induced transcriptional responses, supporting a role in host inflammatory response rather than a validated bacterial-infection biomarker [28]. UBXN7 regulates the balance between NRF2 and HIF-1α under oxidative stress and hypoxia, linking it to metabolic adaptation [29]. A circular RNA generated from the UBXN7 locus has also been associated with diabetic kidney disease, but circUBXN7 is not equivalent to the UBXN7 protein measured here [30]. Accordingly, GTF2I, UBXN7, LIFR and FGF4 remain biologically plausible, potentially novel candidate proteins whose direct comorbidity associations require confirmation.

## Discussion

The main contribution of this study is a disease-contextualized discovery strategy. Instead of assuming that a biomarker separating a comorbidity from healthy individuals will remain informative in COPD, the workflow begins with COPD-centred network hypotheses and tests those hypotheses directly in COPD–comorbidity expression contrasts. This framing matters because systemic inflammation, smoking-related injury and treatment effects can alter the baseline molecular state of COPD patients.

The dual-stage design reduces—but cannot eliminate—the risk of computational false positives. HPN topology and transcriptomic discrimination are generated from different information sources and impose different constraints. A candidate must first be functionally proximate according to the FDM and then display a useful empirical expression pattern. The five-protein panel should nevertheless be understood as a compact set of testable hypotheses, not as a clinical assay. Public-data cross-validation remains internal to the assembled repository, and literature concordance is not a substitute for blinded independent validation.

The revised interpretation also avoids three sources of overstatement. First, the FDM’s 0.811 balanced accuracy is an apparent estimate from training-set predictions, not a held-out accuracy. Second, the historical 224-protein set is supported by the COPD FDM score alone; affected comorbidity-specific CNN columns and the biological interpretations based on them have been removed. Third, the expression stage is cross-sectional. Therefore, the term ‘progression biomarker’ would be premature: a longitudinal design with samples collected before comorbidity development is required to establish temporal prediction.

The individual proteins illustrate how the framework can recover both established and less obvious biology. SRGN provides a positive-control-like example because its tumour-promoting role in NSCLC is experimentally documented. The other four proteins occupy plausible inflammatory, hypoxic, proteostatic or tissue-repair pathways but lack direct validation for the precise clinical contrasts. Their relative novelty is not a defect; it is the hypothesis-generating value of the analysis. Future work should prioritize replication of the expression signal, measurement at the protein level, direction and threshold stability, and mechanistic tests in disease-relevant models.

The framework is portable. Replacing COPD effectors with those of another index disease would preserve the same conceptual sequence: construct a disease-centred network neighbourhood, score functional proximity, test candidates in patient-derived molecular data and retain only findings supported by transparent error estimates. The greatest methodological gain will come from prospectively defining labels, locking feature transformations and evaluating the complete pipeline in an untouched external cohort. An additional potential application of this framework is the investigation of comorbidities for which molecular data from patients with confirmed concurrent diagnoses are scarce or unavailable. By comparing independently characterized disease cohorts against an index-disease molecular background, the approach may generate candidate biomarkers for subsequent prospective evaluation. If validated, this strategy could be particularly useful for studying uncommon comorbidities and rare diseases, where assembling sufficiently large cohorts with multiple concurrent diagnoses is challenging.

## Methodological Considerations and Future Validation

Several limitations should be considered, although the study design incorporates measures intended to reduce their impact. First, the reported FDM metrics were reconstructed from predictions on the training records and therefore represent apparent or resubstitution performance rather than an unbiased estimate of generalization. We consequently use the FDM only as a prioritization mechanism within the network, not as a clinically validated classifier. Moreover, the candidates produced by this stage were subjected to a second, independent source of evidence: their ability to discriminate COPD-associated comorbidity states in patient-derived transcriptomic data. Definitive assessment of FDM generalization will nevertheless require a locked model and a previously untouched test set.

Candidate filtering and final-model selection were reconstructed from archived analysis outputs, which may introduce selection bias, and multiplicity-adjusted inference could not be recovered. To limit overinterpretation, the 224-protein candidate set was defined solely by the reproducible COPD FDM score threshold of 0.50, while the final proteins were retained according to the documented historical cross-validation procedure. Balanced accuracy and MCC were recalculated from the stored predictions because they are invariant to the arbitrary assignment of the positive and negative classes. The resulting proteins are therefore presented as exploratory candidates rather than statistically confirmed biomarkers, and no claims based on uncorrected significance testing are made.

Public transcriptomic cohorts inevitably differ in platform, sampled tissue, preprocessing, annotation depth and clinical composition. To improve comparability without assuming that all GEO records constitute a single homogeneous cohort, transformation and standardization were performed within each source dataset, disease labels were harmonized, provenance was retained and every COPD–comorbidity comparison was analysed separately. These measures reduce scale-related variation and prevent inappropriate global pooling, although they cannot eliminate every batch effect or unmeasured confounder. Replication in prospectively collected cohorts using standardized sampling and measurement protocols will therefore be needed.

The available comparisons are cross-sectional and can demonstrate discrimination between molecular states, but they cannot establish whether a marker predicts the future onset or progression of a comorbidity. We have therefore avoided temporal or causal interpretations and describe the selected proteins as candidate discriminators of COPD-associated comorbidity states. Longitudinal studies in which patients are sampled before and during comorbidity development would be required to establish prognostic value.

The final classifiers deliberately use a single molecular feature and do not adjust for age, sex, smoking, medication, COPD severity or other clinical covariates. This design provides transparent and readily interpretable models and tests whether each protein contains an independent discriminatory signal, but it does not demonstrate incremental value beyond established clinical variables. A subsequent validation phase should combine the selected proteins with relevant covariates and compare clinical-only, molecular-only and integrated models in independent cohorts.

Finally, transcript abundance does not necessarily correspond to protein abundance, biological activity or accessibility in a practical assay. The literature-based contextualization and agreement between network prioritization and patient-derived expression provide orthogonal support for biological plausibility, but they do not replace direct measurement. The five proteins should consequently be regarded as candidates for targeted validation using protein-level assays, followed by prospective evaluation of analytical reproducibility, clinical discrimination and assay feasibility.

## Conclusions

We present a corrected and traceable workflow for discovering candidate biomarkers of COPD-associated comorbidity states. An HPN-based functional-distance model reduced a COPD-centred network search to 224 candidates, and independent patient-derived transcriptomic comparisons nominated GTF2I, UBXN7, LIFR, SRGN and FGF4 for five comorbidity contrasts. The framework—not the assertion of a clinically ready panel—is the principal result. It combines mechanistic network prioritization with empirical molecular discrimination and can be transferred to other index disease–comorbidity systems. The five proteins should now be evaluated in independent, longitudinal, clinically characterized cohorts and at the protein level. The accompanying repository-coverage and disease-effector tables document the patient-derived and network resources underlying these analyses and provide a reusable starting point for extending the framework to additional comorbidities.

## Data and code availability

The analyses were performed using independently developed proprietary computational software, trained model objects, and curated datasets derived from publicly available resources, including the Gene Expression Omnibus (GEO). The underlying software source code and trained model objects are not publicly distributed. The methodological procedures are described in the manuscript, and the data supporting the reported findings are provided in the article or derived from publicly accessible sources. This information is intended to facilitate independent reproduction of the analyses by other researchers using their own computational implementations.

## Author contributions and competing interests

The computational software underlying this study was independently conceived, designed, developed, and implemented by J.M.M.B., drawing upon extensive expertise in computational systems biology, network-based pharmacology, and mechanistic modeling, and using exclusively personal computational resources.

E.R. contributed to the operational guidance, systematic monitoring, and validation of computational workflows and procedures throughout the development process, supporting their consistency, reliability, and methodological robustness.

The software was developed independently, without institutional funding, infrastructure, software licenses, or technical development resources from the Universitat Autònoma de Barcelona (UAB) or any other public or private institution. Its source code is not distributed as part of this publication.

X.D.R. provided scientific supervision and expert guidance in the biological and molecular interpretation of the findings, contributing to their mechanistic contextualization, critical scientific discussion, and assessment of their biological relevance. X.D.R. also contributed to the critical review and intellectual refinement of the manuscript.

The scientific contributions of all authors were instrumental in the interpretation, validation, and presentation of the study’s findings.

## Acknowledgements

Part of this work was presented as a poster by the authors at the 2025 Cold Spring Harbor Meeting on Network Biology, held in Cold Spring Harbor, NY, USA, on March 11–15, 2025. The poster was entitled “Identification of COPD Comorbidity Biomarkers Using Network Biology and Deep Learning, Validated with Real-World Data”.

